# Parvalbumin and somatostatin inhibitory microcircuits perform distinct roles in stereo matching

**DOI:** 10.64898/2026.07.15.738526

**Authors:** Mandi C Severson, Jason M Samonds, Carrie Barr, Nicholas J Priebe

**Affiliations:** Center for Learning and Memory, The University of Texas at Austin, Austin TX USA; Center for Perceptual Systems, The University of Texas at Austin, Austin TX USA; Department of Neuroscience, The University of Texas at Austin, Austin TX USA

## Abstract

We readily perceive stereoscopic depth in correlated random dot stereograms (RDS), in which every dot presented to one eye has a matching dot in the other eye. In contrast, anti-correlated RDS, in which corresponding dots have opposite contrast polarity between the two eyes, do not evoke a perception of depth. Nevertheless, anti-correlated RDS still generate local same-polarity matches within the receptive fields of individual neurons. Inhibitory circuitry has been proposed to suppress these false matches and thereby prevent the perception of illusory depth. To investigate how inhibitory circuits in visual cortex contribute to this process, we measured disparity selectivity to correlated and anti-correlated RDS in two major classes of inhibitory neurons in mice: parvalbumin-expressing (PV+) and somatostatin-expressing (SST+) neurons. We found that the disparity tuning of PV+ neurons was highly correlated with the average activity of surrounding neurons. In contrast, the disparity tuning of SST+ neurons was poorly predicted by local population activity, suggesting that PV+ and SST+ neurons integrate excitatory inputs through distinct mechanisms. Optogenetic suppression of PV+ or SST+ neurons further revealed divergent functional roles. Although both cell types contributed to sharpening selectivity for correlated RDS, only SST+ neurons reduced selectivity for anti-correlated RDS. These findings suggest that PV+ and SST+ circuits both enhance responses to correct binocular matches in correlated RDS, whereas SST+ circuits contribute to suppressing false matches in anti-correlated RDS.

## Introduction

Depth can be computed from disparities between the left and right retinotopic projections, but the brain must determine which points from one retinal image correspond to the same physical points in the other retinal image, known as the stereo correspondence problem. One fundamental challenge of this computation is that there are many potential interpretations, and yet we infer depth from stereo information seamlessly. Identifying false matches, or the incorrect matching of features from the left and right eyes, is a key strategy to limit the set of plausible interpretations.

Our understanding of binocular integration in the primary visual cortex (V1) has been guided by the disparity energy model (Ohzawa et al., 1990). In this model, binocular signals are initially combined by polarity-sensitive simple cells, which converge onto complex cells that encode disparity independent of polarity. To avoid potential monocular artifacts, experimentalists use random dot stereograms (RDS) to measure neural responses to disparity. In RDS, changes in disparity are not associated with discernible changes in monocular images (Julesz, 1969). The disparity energy model does well at explaining disparity tuning to RDS for complex cells in V1 (Prince et al., 2002a; Prince et al., 2002b). Because the disparity information is initially generated by polarity sensitive simple cells, the energy model predicts that disparity tuning to stereo matches of opposite polarity (matching black with white dots or “anti-correlated RDS”) are inverted relative to “correlated” RDS. Anti-correlated RDS generally do not produce the perception of depth like standard correlated RDS (Julesz, 1969; Julesz and Tyler, 1976; Cogan et al., 1993; Cumming et al., 1998; Hibbard et al., 2014), though neurons in the primary visual cortex (V1) respond to anti-correlated RDS in a way similar as the model (Cumming and Parker, 1997; Ohzawa et al., 1997; Lippert and Wagner, 2001; Nieder and Wagner, 2001; Samonds et al., 2013; La Chioma et al., 2020). The peaks in an anti-correlated RDS disparity tuning curve represent local false matches. Therefore, it is possible to probe the responses to false matches by presenting anti-correlated RDS, thus offering a method to examine how the visual system rejects these matches.

One important difference between the disparity energy model’s predictions and neurophysiological data is that the magnitude of disparity induced modulation is smaller for anti-correlated than correlated RDS (Cumming and Parker, 1997; Ohzawa et al., 1997). The disparity energy model predicts that these would be the same magnitude (Ohzawa et al., 1990). This suggests that V1 is at least partially solving the stereo correspondence problem and represents an intermediate stage in the process of suppressing false matches in anti-correlated RDS (Samonds et al., 2013).

Several hypotheses have been proposed about how the populations of neurons in the visual cortex could suppress false matches. First, coarse-to-fine processing of disparity, with coarse processing preceding and constraining fine processing (Menz and Freeman, 2003; Chen and Qian, 2004). Second, recurrent interactions (both suppressive and facilitative) among disparity-tuned neurons could refine disparity tuning (Samonds et al., 2009; Samonds et al., 2013). Third, phase-disparity neurons could act as “lie detectors” to signal false matches (Read and Cumming, 2007). Fourth, the alignment of receptive fields in V1 in a push-pull manner could reduce the response to false matches (Tanabe et al., 2011). Lastly, thresholding could effectively suppress false matches (Lippert and Wagner, 2001). One common element in many of these hypotheses is that they predict that inhibitory neurons would play a significant role in suppressing false matches.

To ascertain how cortical inhibitory neurons might contribute to solving the stereo correspondence problem, we measured disparity selectivity of two major subclasses of inhibitory neurons, parvalbumin-expressing (PV+) and somatostatin-expressing (SST+) in mouse binocular visual cortex. We also measured the impact of suppressing each of the inhibitory cell classes on the disparity selective responses of the remaining neuronal population. Our results demonstrate that both PV+ and SST+ inhibitory neurons enhance correct matches in correlated RDS, but only SST+ neurons appear to play a role in suppressing false matches that occur in anti-correlated RDS.

## Materials and Methods

### Preparation of animals

The University of Texas at Austin Institutional Animal Care and Use Committee approved all procedures, and they are in accordance with the National Institutes of Health Guide for the Care and Use of Laboratory Animals. For imaging, we crossed tetO-GCaMP6s mice (Jackson Labs, # 024742) with CamKII-tTA mice (Jackson Labs, # 007004) to express the GCaMP6s calcium indicator in excitatory neurons in 10 mice (3 females, 7 males) (Wekselblatt et al., 2016). We additionally imaged 3 GCaMP-Thy1 animals (2 females, 1 male) (Jackson Labs, # 025393), which express GCaMP6f in excitatory neurons under the Thy1 promoter (Dana et al., 2014). To identify inhibitory neurons, we used 3 PV-Ai14 and 10 PV-Cre mice (8 females, 5 males) (Scholl et al., 2015) and 13 SST-Cre mice (7 females, 6 males) (Lovett-Barron et al., 2014). For the optogenetics experiments, we used an additional 9 PV-Cre (5 females, 4 males), 8 SST-Cre (2 females, 6 males), and 2 GCaMP6s (2 males) mice. For each set of experiments, mice were imaged using two-photon microscopy when they were between 3-7 months of age.

In preparation for two-photon microscopy experiments, a titanium bar was secured to the skull using dental acrylic as described previously (Samonds et al., 2019). Animals were anesthetized during this procedure using isoflurane anesthesia (1-3%). After titanium head bars were placed, a 3 or 4 mm craniotomy was made over the binocular region of V1 and a glass window was placed over top and secured with cyanoacrylate. If the animals were PV-Cre or SST-Cre, they also received adeno-associate virus (AAV) injections before the cranial window was placed.

Viral mixes were chosen to express GCaMP6f in all neurons (hSYNCMV GCamp6f TPI I6 AAV) and TdTomato in neurons expressing Cre-recombinase, either PV+ or SST+ neurons (mSYN-422 CMV tetO(4) nls-tdTomato (lox) AAV). Viral mixes for the optogenetics experiments also included Archaerhodopsin-T (Arch-T) to express in neurons with Cre-recombinase (h56D CMV tetO(4) ArchT tomato targeting (lox) AAV). Mice were anesthetized with inhaled isoflurane (1%–3% in oxygen), and body temperature was maintained at 37° C. Injections were performed using a stereotaxic apparatus (Sutter Instruments and Schaublin) fitted with a Nanoject II microinjector (Drummond Scientific and World Precision Instruments). Pulled-glass pipettes back-filled with mineral oil were used to deposit virus mixes. 75 nL virus was deposited in each site in three 25-nl pulses. There were 10-15 injection sites per animal. Animals were given 4 weeks for the virus to fully express before imaging.

### Retinotopic mapping

Fluorescence was measured optically across the entire imaging window using the previously described intrinsic imaging and stimulation setup (Kalatsky and Stryker, 2003; Nauhaus and Ringach, 2007; Garrett et al., 2014). Mice were head-fixed on a custom stand and lightly anesthetized with 0.5% isoflurane, and silicone oil was applied to their eyes. A contrasting checkered bar was used as a visual stimulus, presented for twelve trials and alternating randomly between the four cardinal directions. Stimuli were projected onto a 64 x 46-cm screen, placed at a 30-degree angle, 10 cm from and centered on the right eye, using an LC4500-UV projector (Keynotes Photonics). A pco.panda 4.2 sCOMAS camera (Excelitas) equipped with a 25 mm C VIS-NIR fixed focal length lens (Edmond Optics) and a 525 nm fluorescence filter (Thor Labs) was used for imaging. For illumination, an XCite 110 LED illuminator (Excelitas) with a 460 nm filter (Thor Labs) was used. External light was blocked between the imaging window and the lens with clay. Using the retinotopic maps obtained from these experiments, we targeted binocular visual cortex in the center of the visual field.

### Two-photon microscopy

We measured changes in fluorescence from the calcium indicator GCaMP6s, which was expressed in excitatory neurons of CamKII-tTA/tetO-GCaMP6s mice (Wekselblatt et al., 2016). Our methods were similar to those previously described (Samonds et al., 2019). We presented 6-second dynamic random dot stereograms (DRDS; 6 Hz, 7.4° dots, 200 black and 200 white dots) interleaved with 6-second periods of a mean gray screen. A circular polarization alternator (DepthQ/Lightspeed Design) was synchronized with an Optoma HD27 projector (120 Hz refresh rate) to rear-project images onto an RP3D polarization-preserving screen (Severtson Screens) placed 22 cm from the mouse. Stereoscopic depth was created by shifting dots to each eye every other frame, with passive circular polarization filters placed in front of the mouse’s eyes. The dynamic random dot stereograms (DRDS) were 70° wide and 100° high, and binocular disparity was varied at 0°, ±1.85°, ±5.55°, and ±9.20° within the vertical central 65% of the display, while remaining at 0° on the sides. Ten different DRDS were used for each disparity. For anti-correlated DRDS, each black dot in one eye was matched with a white dot and vice versa. Imaging was performed using a custom-built two-photon resonant mirror scanning microscope with a mode-locked 920-nm Chameleon Ultra Ti:Sapphire laser (Coherent Technologies) (Scholl et al., 2015). A 16× water objective (0.8 numerical aperture; Nikon) focused green light, collected from a 0.4 to 0.5 mm-square region by photomultiplier tubes. Images (256 × 455 pixels) were acquired at a frame rate of 30.9 Hz using custom software (Labview; National Instruments), with focal planes ranging from 130 to 200 μm below the cortical surface. The objective was oriented normal to the cortical surface, and clay was used to block external light. During imaging, we also tracked the pupils of both eyes using a previously described method (Samonds et al., 2018; Samonds et al., 2019; Brown et al., 2025). Two infrared cameras (30 frames per second) were mounted perpendicular to each orbital axis while the mouse was head-fixed and running on a floating trackball (Dombeck et al., 2007). Pupil position was tracked using DeepLabCut, and calibrated to degrees of visual angle based on a similarly sized artificial eyeball with known rotations.

### Two-photon data analysis

We analyzed images using custom Matlab software. Initially, we bandpass filtered the images over time by applying an 8-frame moving average and subtracting a 1000-frame median filter. Neurons were manually identified in images and video based on changes in brightness, size, and shape. We drew masks around identified neurons, discarding any overlapping regions. The neuropil was defined as all pixels outside these masks. For each neuronal mask, a neuropil annulus was created, 4 pixels wide and 3 pixels away along the 455-pixel dimension, and 2 pixels wide and 2 pixels away along the 256-pixel dimension, with any overlap with other masks discarded. We then computed the mean brightness of pixels within each mask, the neuropil, and neuropil annuli. To derive ΔF, we subtracted the average intensity calculated 4 seconds before all stimuli from these values. We then divided these differences by the median intensity over the entire experiment to obtain ΔF/F. For masks, we further subtracted the corresponding neuropil annulus value for a final neuropil-corrected ΔF/F.

Visually responsive neurons were identified as those with a significantly greater average response during the 6-second stimulus period compared to the 4 seconds pre-stimulus (sign test, p < 0.05, n = 70 trials). Disparity-tuned neurons showed a significantly different average response with respect to disparity (Kruskal-Wallis test, p < 0.05, n = 10 trials, 7 disparities). The disparity selectivity index (DSI) was calculated using the resultant vector (Ringach et al., 2002):

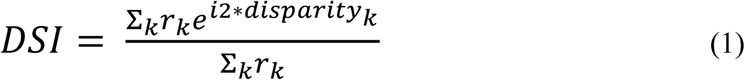

Here, *r*_k_ represents the response to each disparity. We assumed tested disparities wrapped around at their extremes for this circular calculation. DSI is 0 if all responses are equal and 1 if a neuron responds to only a single disparity. We fit a Gabor function to the average responses *f* across the 7 disparities *d*, but this fit was not used for analysis:

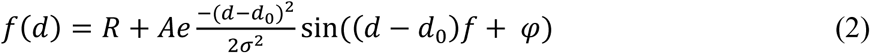

Pearson’s correlation was computed between the 7 mean disparity responses to correlated and anti-correlated RDS (r_correlated-anti-correlated_). Similarly, we used Pearson’s correlation to compute the correlation between disparity tuning to correlated or anti-correlated RDS for a single neuron and the average tuning for all significant visually responsive neurons within a radius of 125 µm of that neuron. The average tuning for the surround population was computed by first normalizing each tuning curve by the average preferred disparity response to correlated RDS. We only included neurons with at least 5 surrounding visually responsive neurons. We also used Pearson’s correlation to compute signal correlation using the 7 mean disparity responses for a pair of neurons. And finally, we used Pearson’s correlation to compute the noise correlation between a pair of neurons. To compute noise correlation, we used all 70 paired responses and z-scored each response based on the mean and standard deviation of the 10 responses to each of the 7 disparities.

### Optogenetics

We used a light-emitting diode (LED) (CoolLED) delivering 590 nm light routed through the objective to induce optogenetic suppression of PV+ or SST+ neurons. In order to reduce bleedthrough, we added a pi cell liquid crystal shutter with a compensating polarizer (Bolder Vision Optik) in front of the photomultiplier tube (PMT). The LED was synchronized with the pi cell and only delivered light when the shutter was closed. Analysis was performed on the frames in which the shutter was open and LED was off. LED power was calibrated by measuring it at the objective. The LED was turned on 2 s before stimulus offset and turned off 0.5 s after stimulus offset. Neurons with a significant decrease (t-test, p < 0.05) in response when the LED was on in the 2 seconds before stimulus onset were classified as PV+ or SST+ and all other neurons were classified as PV- or SST-.

### Statistics

All statistical analyses were performed using Matlab, with specific test details provided in the relevant figure legends. While we did not use a formal power analysis to determine sample size, our numbers of animals, trials, and neurons are comparable to or larger than those in previous studies (Scholl et al., 2015; Samonds et al., 2019). Data collection and analysis were not blinded to subject conditions. No assumptions were made about the underlying distributions and all statistical tests for behavioral and neuronal data were non-parametric, including Kruskal-Wallis, Wilcoxon rank-sum, and sign tests.

## Results

We examined the roles of two inhibitory cell classes, PV+ and SST+ inhibitory neurons, in processing binocular information in V1. Both the inputs of these inhibitory cells and their targets differ: PV+ inhibitory neurons receive strong feedforward excitatory input from layer 4 and recurrent connections from layer 2/3, while excitatory input to SST+ inhibitory neurons comes mainly from layer 2/3 (Dantzker and Callaway, 2000; Xu and Callaway, 2009; Adesnik et al., 2012); PV+ inhibitory neurons are known to synapse onto the cell bodies of excitatory neurons (Jones and Hendry, 1984), while SST+ inhibitory neurons target the dendrites (Kawaguchi and Kubota, 1997; Wang et al., 2004; Xu and Callaway, 2009). The differences suggest that these cell classes could play distinct roles in shaping responses to correlated and anti-correlated RDS. We first examined the disparity sensitivity of excitatory neurons before measuring and manipulating these inhibitory cell classes.

### Excitatory neurons in mice respond to disparity in anti-correlated RDS

Similar to macaques (Prince et al., 2002a; Prince et al., 2002b) and cats (Ohzawa et al., 1990; Deangelis et al., 1991; Ohzawa et al., 1997), mice have disparity-tuned neurons in V1 to RDS with responses that are well described by Gabor functions, albeit for much larger disparities (La Chioma et al., 2019; Samonds et al., 2019). As the disparity energy model predicts (Ohzawa et al., 1990; Cumming and Parker, 1997), binocular neurons in mouse V1 also respond to anti-correlated RDS with responses described well by a Gabor function that are on average 180 degrees out of phase with respect to correlated RDS (La Chioma et al., 2020). Similar to macaques (Cumming and Parker, 1997), the disparity amplitude modulation (the maximum – minimum response) for anti-correlated RDS for mice is smaller than predicted by the disparity energy model (La Chioma et al., 2020).

We used two-photon microscopy to measure visually-evoked responses in transgenic mice that express GCaMP6s in excitatory neurons (Wekselblatt et al., 2016). Within the box (Figure 1A), we show a representative example of the significant visually-responsive (gray) and disparity-tuned neurons (black). These include neurons that responded most to crossed disparities projected in front of our screen (Figure 1B, “tuned near” or rendered depths less than 22 cm), most to zero disparities (Figure 1B, “tuned zero” or rendered depths near 22 cm), and most to uncrossed disparities projected behind our screen (Figure 1B, “tuned far” or rendered depths greater than 22 cm). The solid black points and lines represent disparity-dependent responses to correlated RDS and the open circles and dashed lines represent disparity-dependent responses to anti-correlated RDS (Figure 1B). The tuning to anti-correlated RDS was generally 180 degrees out of phase (or inverted) and had smaller amplitude modulation compared to correlated RDS in many of these examples. Similarly, across all our data, the distribution of the correlation between correlated and anti-correlated RDS tuning curves was skewed towards negative one and consistently below zero for individual mice, suggesting that it was generally inverted (Figure 1C). Most response amplitude modulations for neurons fell below the principal diagonal and the amplitude modulation ratio (AMR, amplitude modulation for anti-correlated RDS divided by amplitude modulation for correlated RDS) was consistently below one for individual mice (Figure 1D). These results are consistent with those measured by La Chioma et al. (2020) and similar to what we observe when we applied the same metrics to electrophysiological spike rates measured from V1 neurons in the macaque (Figure S1; Samonds et al. 2013). These data show that false match responses to anti-correlated RDS are partially suppressed in mouse visual cortex with a magnitude similar to V1 in non-human primates.

**Figure 1.**
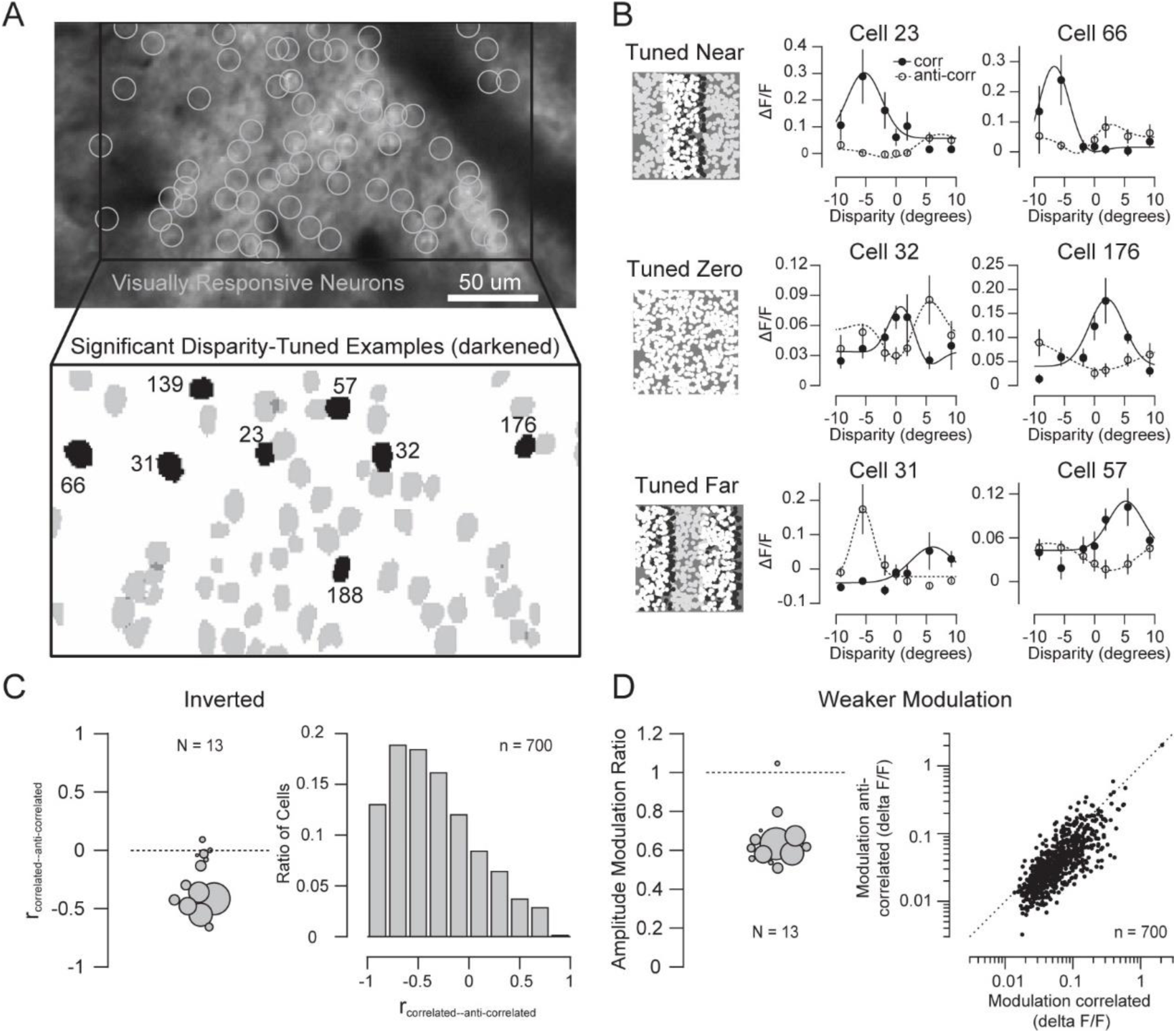
Disparity tuning to anti-correlated stereograms is inverted and has weaker modulation with respect to correlated stereograms. **A**, Responses of excitatory neurons in transgenic mice expressing GCAMP6s were measured with two-photon microscopy. First, neurons were identified with significant visual responses (sign test, *p* < 0.05, *n* = 70 trials) and significant disparity tuning (Kruskal-Wallis test, *p* < 0.05, *n* = 10 trials, 7 disparities). **B**, Examples of disparity tuning to correlated (corr) dynamic random dot stereograms (DRDS) and anti-correlated (anticorr) DRDS fit with Gabor functions. Error bars are standard error. **C**, The correlation between corr and anticorr disparity tuning was significantly negative across mice (Wilcoxon signed rank test, *p* = 0.003) and all neurons (Wilcoxon signed rank test, *p* < 0.001), suggesting that anticorr tuning is inverted with respect to corr disparity tuning. Area of circles for individual mice represent the number of neurons. **D**, The modulation (maximum-minimum response) for anticorr disparity tuning was significantly smaller than for corr disparity tuning across mice (Wilcoxon signed rank test, *p* < 0.001) and all neurons (Wilcoxon signed rank test, *p* < 0.001). The amplitude modulation ratio for individual mice is anticorr modulation divided by corr modulation.

### How inhibitory neurons could suppress responses to false matches

One potential mechanism for selectively suppressing responses to false matches is for inhibitory neurons to integrate excitatory responses in a disparity-dependent manner. When neurons are aligned by their preferred disparity or peak response to correlated RDS, the peaks of their responses to anti-correlated RDS do not align (Samonds et al., 2013). We illustrate this using four example excitatory neurons from Figure 1A. Aligning disparity-tuned excitatory neurons with a preferred disparity of –5.5° reveals consistent tuning to correlated RDS (Figure 2A, left), whereas their preferred disparities for anti-correlated RDS are scattered (Figure 2A, right). To quantify this effect, we aligned all 700 excitatory neurons by their preferred disparity and normalized each neuron by its response at the preferred disparity. Averaging across the population produced a much flatter tuning curve for anti-correlated RDS than for correlated RDS (Figure 2B), resulting in an AMR substantially smaller than what we observed in individual neurons (Figure 1D). Thus, selective integration based on correlated-RDS disparity on average reduces the magnitude of the negative correlation (inversion) between correlated and anti-correlated tuning and attenuates disparity-dependent modulation to anti-correlated stimuli. If inhibitory neurons selectively integrate excitatory inputs with similar disparity preferences, they would reduce modulation to false matches in their responses and could in turn suppress excitatory responses to false matches in anti-correlated RDS.

**Figure 2.**
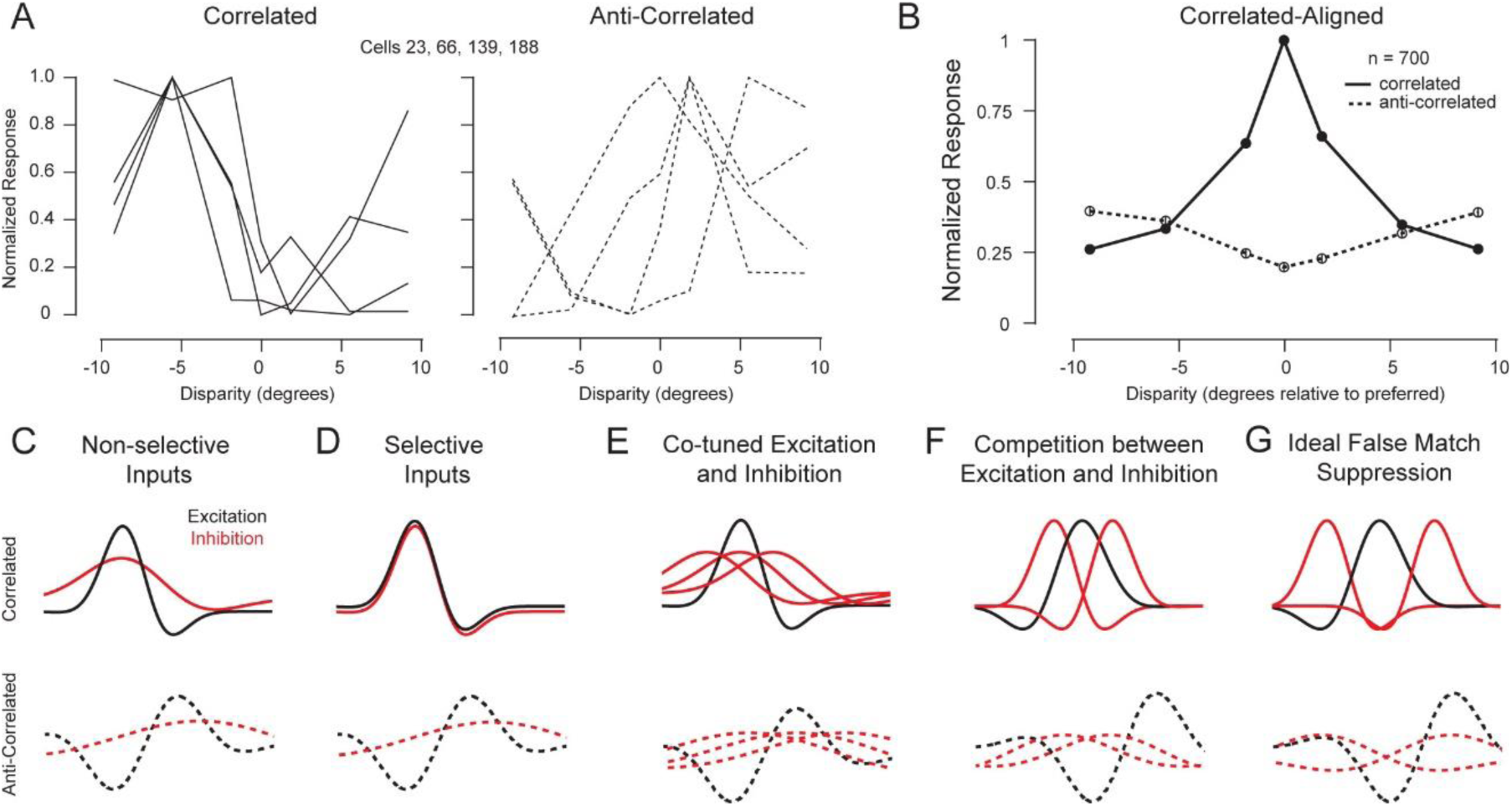
Potential mechanisms of suppressing false matches with inhibitory neurons. **A**, Four example excitatory neurons with similar disparity tuning to correlated RDS (left) and their corresponding disparity tuning to anti-correlated RDS (right). **B**, Population average of normalized tuning curves for correlated and anti-correlated RDS both aligned to the preferred disparity to correlated RDS. **C**, Predicted disparity tuning curve for an inhibitory neuron that non-selectively integrates from surrounding excitatory neurons. **D**, Predicted disparity tuning curve for an inhibitory neuron that selectively integrates from surrounding excitatory neurons with similar disparity tuning to correlated RDS. **E**, Alignment of excitatory and inhibitory disparity tuning curves for inhibition with matched disparity tuning to excitatory targets on average. **F**, Alignment of excitatory and inhibitory disparity tuning curves for inhibition that targets non-preferred disparities for excitatory neurons on average. **G**, An ideal false match suppressing circuit where inhibitory neurons target non-preferred disparities in excitatory neurons and have positive correlation between correlated and anti-correlated RDS disparity tuning.

Because disparity is represented heterogeneously in mouse visual cortex, the disparity selectivity of inhibitory neurons provides insight into how they integrate excitatory inputs (Scholl et al., 2015). An inhibitory neuron that pools indiscriminately from local excitatory neurons would be expected to exhibit weak or flat disparity tuning. However, the coarse organization of preferred disparity in mouse visual cortex (La Chioma et al., 2019; Brown et al., 2025) creates local biases that produce broad tuning reflecting the average activity of nearby excitatory neurons (Scholl et al., 2015) (Figure 2C). Because the anti-correlated tuning peaks of neighboring excitatory neurons are misaligned (Figure 2B), such non-selective integration should yield relatively flat anti-correlated tuning curves compared with correlated tuning (Figure 2C). In contrast, an inhibitory neuron that selectively integrates excitatory inputs with similar disparity preferences should exhibit tuning selectivity comparable to that of individual excitatory neurons, and its tuning should deviate from that predicted by non-selective local pooling (Figure 2D). Such neurons would most closely match the population-based prediction shown in Figure 2B and would exhibit an even greater departure from perfectly inverted tuning between correlated and anti-correlated RDS (Figure 2D).

Disparity-tuned inhibitory neurons could suppress disparity-tuned excitatory neurons with multiple potential strategies. Excitatory and inhibitory inputs to excitatory neurons might be co-tuned for the same preferred disparity for correlated RDS (Figure 2E). If the inhibition is modest and untuned or broadly-tuned compared to excitatory neurons, inhibition generally reduces the responses without changing the shape of the tuning curve by lowering the gain (Atallah et al., 2012; Wilson et al., 2012). If the inhibition is strong and broadly-tuned, it will reduce some responses below threshold and effectively sharpen tuning (Lee et al., 2012). It is unclear how inhibition would reduce responses to false matches with this organization, since inhibition would have a similar effect of reducing modulation for correlated and anti-correlated disparity tuning. Alternatively, inhibitory neurons may selectively target excitatory neurons at their non-preferred disparities (Figure 2F). In this case, the excitatory and inhibitory inputs would be competing with differing preferred disparities. With narrow or broad tuning, inhibition would sharpen disparity tuning for correlated RDS (Figure 2F, top). The scattering of anti-correlated disparity tuning curves results in weaker sharpening or even eliminates sharpening of disparity tuning for anti-correlated RDS (Figure 2F, bottom). In the most extreme case, if the scattering of anti-correlated RDS peak responses results in positive correlation between disparity tuning to correlated and anti-correlated RDS, inhibition would sharpen disparity tuning for correlated RDS and flatten tuning for anti-correlated RDS (Figure 2G). This would be the ideal inhibitory neuron for suppressing false matches.

To understand how PV+ or SST+ inhibitory neurons might suppress false matches, we present analyses and experiments to (1) measure disparity tuning in inhibitory subclasses, (2) compare disparity tuning between correlated and anti-correlated RDS in inhibitory subclasses, (3) characterize how inhibitory neurons integrate excitatory inputs, (4) characterize how inhibitory and excitatory neurons are connected with respect to disparity tuning, and (5) characterize how inhibitory neurons suppress excitatory responses. These experiments and analyses demonstrate that PV+ and SST+ neurons play distinct roles in processing stereo information.

### Inhibitory subclasses deviate from the disparity energy model

If inhibitory neurons play a role in suppressing false matches, they may express distinct selectivity for disparity stimuli than excitatory neurons. Previous work has shown that PV+ inhibitory neurons have broad disparity tuning when using sine wave gratings (Scholl et al., 2015), and we initially sought to characterize that selectivity using correlated RDS stimuli. We used two-photon microscopy to measure visually evoked responses in PV-Cre and SST-Cre transgenic mice following virus injection to express GCaMP6s in all neurons (Figure 3A, green) and TdTomato in PV+ and SST+ inhibitory neurons (Figure 3A, red), respectively. The majority of PV-/SST- neurons are represented by excitatory neurons, but they also include a percentage of other inhibitory neuron types, since GCAMP6s was expressed in all neurons. We found that both PV+ and SST+ inhibitory neurons exhibited robust disparity tuning (e.g., Figure 3C and 3F). We found that PV+ inhibitory neuron disparity selectivity for correlated stimuli was weaker compared to PV- neuron disparity tuning though (Figure 3B-D; see also Scholl et al., 2015). Note that although this PV+ neuron is clearly disparity-tuned, it responded robustly to all disparities tested (Figure 3C, all responses above the dashed line). On the other hand, we measured no significant difference for disparity selectivity between SST+ inhibitory and SST- neurons (Figure 3E-G). For SST+, SST- and PV- example neurons, we see that they only responded to some of the disparities tested (Figure 3B, 3E, and 3F, responses above the dashed line). These two measures of disparity selectivity suggest that the roles these two inhibitory cells play in shaping disparity selectivity in excitatory neurons may be distinct.

**Figure 3.**
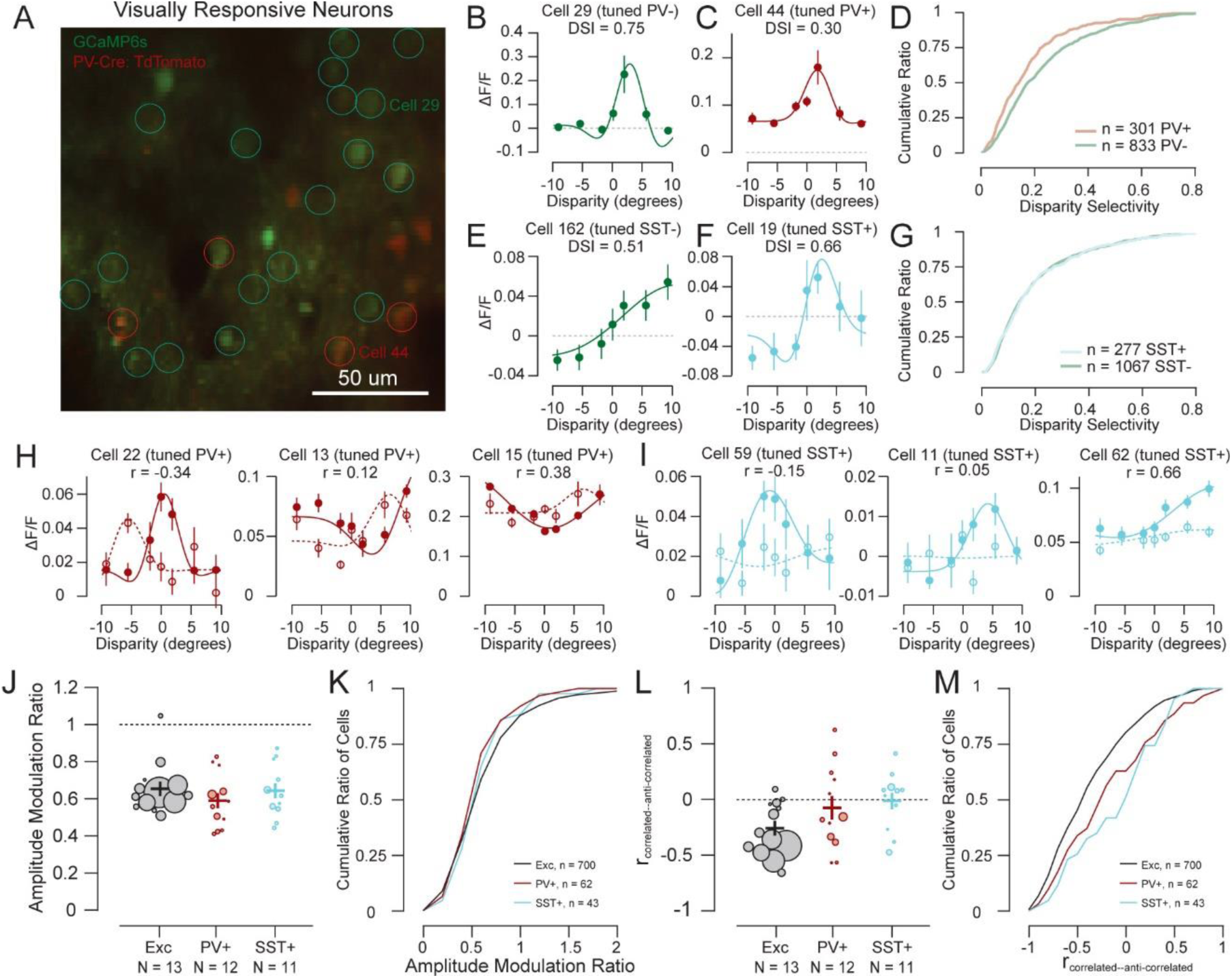
Inhibitory subclasses are selective to disparity, but deviate more from the energy model than excitatory neurons. **A**, Responses of neurons in PV-Cre transgenic mice were measured following virus injection to express GCaMP6s in all neurons and TdTomato in parvalbumin-expressing (PV+) neurons. Visually responsive and disparity-tuned neurons were identified based on changes in green calcium-dependent fluorescence and PV+ neurons were identified based on red fluorescence. **B**, Example disparity tuning curve for a PV- neuron that responded to only a few disparities. **C**, Example disparity tuning curve for PV+ inhibitory neuron that responded to all disparities. **D**, Disparity selectivity was significantly weaker for PV+ versus PV- (left, Wilcoxon rank sum test, *p* < 0.001). **E**, Example disparity tuning curve for a SST- neuron that responded to only a few disparities. **F**, Example disparity tuning curve for a SST- inhibitory neuron that also responded to only a few disparities. **G**, There was no significant difference in disparity selectivity between SST+ and SST- (right, Wilcoxon rank sum test, *p* = 0.98). **H**, Example disparity tuning curves for correlated and anti-correlated RDS for PV+ neurons. I, Example disparity tuning curves for correlated and anti-correlated RDS for SST+ neurons. **J**, Across mice, there is no significant difference in the modulation amplitude between anticorr and corr disparity tuning for PV+ and SST+ neurons compared excitatory neurons (Wilcoxon rank sum test, *p* = 0.29 and *p* = 0.91, respectively). **K**, For all disparity-tuned neurons, there is no significant difference in the modulation amplitude between anticorr and corr disparity tuning for PV+ and SST+ neurons compared excitatory neurons (Wilcoxon rank sum test, *p* = 0.23 and *p* = 0.63, respectively). **L**, Across mice, anticorr disparity tuning is not as negatively correlated with corr disparity tuning for SST+ neurons compared to excitatory neurons (Wilcoxon rank sum test, p = 0.02). **M**, For all disparity-tuned neurons, anticorr disparity tuning is not as negatively correlated with corr disparity tuning for both PV+ and SST+ neurons compared to excitatory neurons (Wilcoxon rank sum test, *p* = 0.005 and *p* < 0.001, respectively).

To cancel false matches prominent in anti-correlated RDS effectively, inhibitory neurons should integrate from excitatory neurons in a disparity-dependent manner to weaken modulation to anti-correlated disparity (Figure 2B). We found that PV+ (e.g., Figure 3H) and SST+ (e.g., Figure 3I) inhibitory neurons responded selectively to disparity in anti-correlated RDS, but the amplitude modulation ratio in PV+ and SST+ neurons was not significantly reduced beyond what was seen in excitatory neurons (Figure 3J-K). The inversion in disparity tuning though differed substantially. Instead of a prominent negative correlation between correlated and anti-correlated tuning curves, the relationship between these tuning curves for PV+ and SST+ neurons was closer to zero (Figure 3L-M). This scatter in the relationship between correlated and anti-correlated disparity tuning should result in flattened disparity tuning to anti-correlated RDS when multiple inhibitory neurons target excitatory neurons. Overall, this suggests that the integration performed by inhibitory neurons results in disparity tuning properties that deviate even farther from the energy model than what we observe in excitatory neurons (e.g., Figure 2B). There were some neurons with positive correlation between correlated and anti-correlated RDS that would potentially correspond to ideal false match suppression (Figure 2G). Specific examples of high correlation generally had relatively flat tuning to anti-correlated disparity though (e.g., Figure 3I, right) and in general, tuning to anti-correlated RDS exhibited unexpected phase shifts with an average correlation close to zero. Nonetheless, these SST+ inhibitory neurons could potentially be effective in suppressing false matches.

### PV+ and SST+ inhibitory neurons integrate disparity information differently across cortical space

The origin of these differences in selectivity for disparity signals between excitatory, PV+ and SST+ neurons likely lies in how these neurons integrate excitatory input. Previous work has shown that PV+ inhibitory neurons have broad disparity tuning due to the nonspecific pooling of nearby excitatory neurons (Scholl et al., 2015). To probe for a circuit difference, we examined the relationship between inhibitory neurons and their surrounding network. Figure 4A shows an example PV+ inhibitory neuron (red) and several example disparity-tuned PV- neurons (green) that are within 125 um of the PV+ inhibitory neuron. Summing the disparity tuning of each of these PV- neurons (Figure 4B) and all other surround visually responsive neurons (Figure 4A) roughly matches the tuning of the neighboring PV+ inhibitory neuron (Figure 4C, solid red versus solid light green).

**Figure 4.**
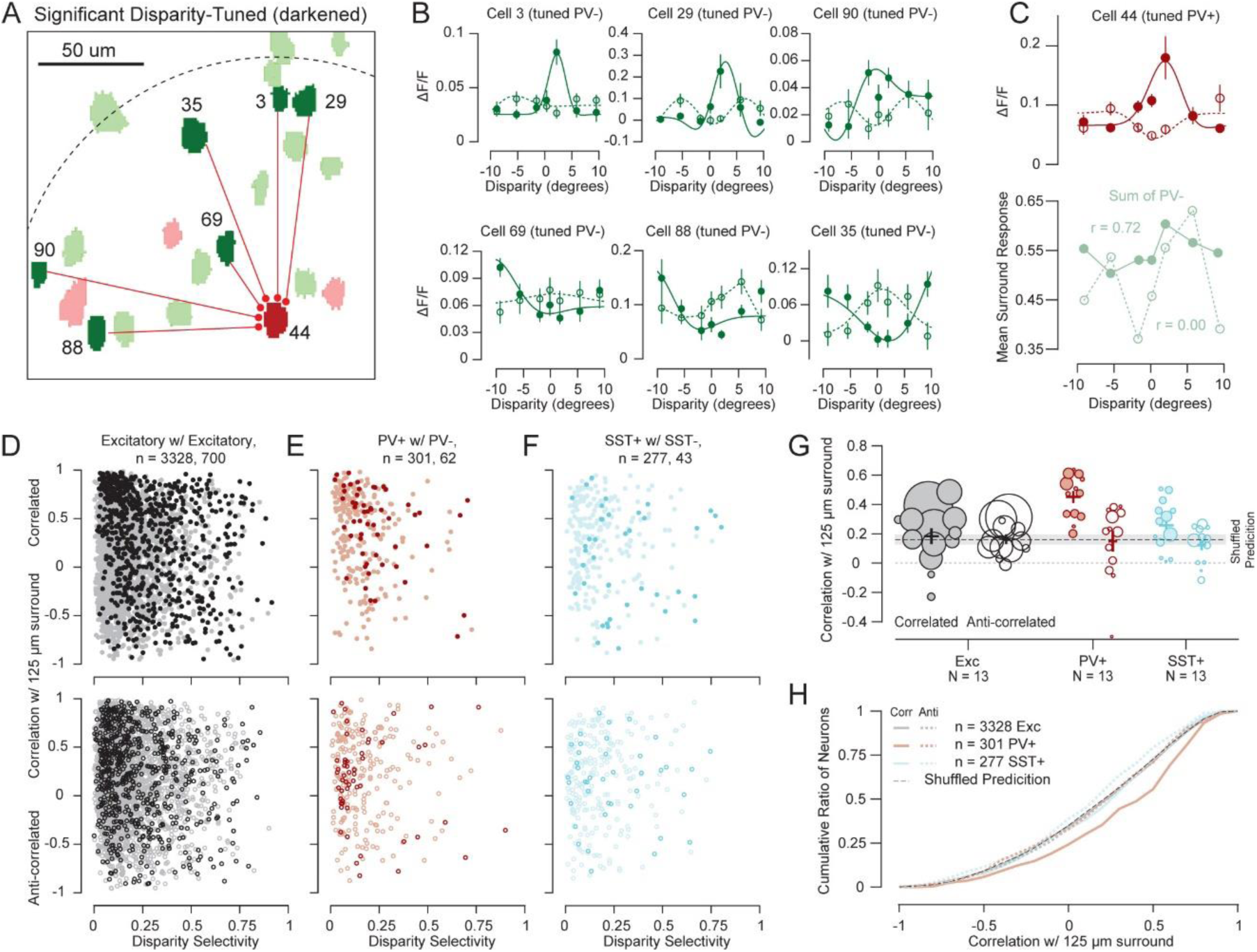
Local integration differs between PV+ and SST+ inhibitory neurons. **A**, Visually responsive (light) and disparity-tuned neurons (dark) were identified and ROIs are noted as green for PV- and red for PV+. **B**, Examples of disparity tuning in PV- neurons to correlated RDS (solid lines, filled circles) and anti-correlated RDS (dashed line, open circles) fit with Gabor functions. Error bars are standard error. **C**, Example of disparity tuning in a PV+ neuron (top) compared to the sum of responses to all PV-visually responsive neurons within a radius of 125 µm (bottom). **D**, The correlation between disparity tuning for a single excitatory neuron and the average of normalized disparity tuning curves for all visually responsive excitatory neurons within a radius of 125 µm. Darkened data points are neurons with significant disparity tuning. Top row is disparity tuning for correlated RDS and bottom row is disparity tuning for anti-correlated RDS. **E**, The correlation between disparity tuning for a single PV+ neuron and the average of normalized disparity tuning curves for all visually responsive PV- neurons within a radius of 125 µm. **F**, Same as E, but for SST-Cre mice where TdTomato was expressed in somatostatin expressing (SST+) neurons. **G**, Across mice, the average correlation within a radius of 125 µm for PV+ neurons for correlated RDS (filled circles) was significantly higher compared to excitatory, SST+ neurons, and the shuffled prediction (Wilcoxon rank sum test for all subsequent comparisons, p = 0.001, p = 0.01, and p < 0.001, respectively). There was no significant difference in local correlation between mice for correlated RDS for excitatory, SST+ neurons, and the shuffled prediction (p = 0.41, p = 0.44, and p = 0.12, respectively). The correlation significantly dropped for PV+ and SST+ from correlated to anti-correlated RDS (p = 0.003 and p = 0.03, respectively), but not for excitatory neurons (p = 0.44). There was no significant difference in local correlation between mice for anti-correlated RDS for excitatory, PV+ neurons, and the shuffled prediction (p = 0.64, p = 1, and p = 0.64, respectively). **H**, These comparisons were even clearer for aggregated data. The average correlation within a radius of 125 µm for PV+ neurons for correlated RDS (solid lines) was significantly higher compared to excitatory, SST+ neurons, and the shuffled prediction (p < 0.001). There was no significant difference in local correlation among neurons for correlated RDS for excitatory, SST+ neurons, and the shuffled prediction (p = 0.79, p = 0.39, and p = 0.55, respectively). The correlation significantly dropped for PV+ and SST+ from correlated to anti-correlated RDS (p < 0.001 and p = 0.05, respectively), but not for excitatory neurons (p = 0.77). The average correlation within a radius of 125 µm for SST+ neurons for anti-correlated RDS (dashed lines) was significantly lower compared to excitatory neurons and the shuffled prediction (p = 0.01 and p = 0.05, respectively). There was no significant difference in local correlation among neurons for anti-correlated RDS for excitatory, PV+ neurons, and the shuffled prediction (Wilcoxon rank sum test, p = 0.69, p = 0.24, and p = 0.97, respectively).

As shown in Figure 2A-B, preferred disparities for anti-correlated RDS do not necessarily line up for neurons with similar preferred disparity for correlated RDS. Therefore, even though disparity tuning for correlated RDS for a PV+ inhibitory neuron can be predicted by surrounding PV- neurons, disparity tuning for anti-correlated RDS might not be as predictable. The anti-correlated disparity tuning of our example PV+ neurons does not match up with the sum of the anti-correlated disparity tuning of the surrounding PV- neurons (Figure 4C, dashed red versus dashed light green).

We quantified this behavior by measuring the correlation between the disparity tuning curve of the inhibitory neuron (Figure 4C, red) and the average of the peak normalized disparity tuning curves of all significantly visually responsive neurons within 125 µm of the inhibitory neuron (Figure 4C, light green). For our example neuron, this correlation was 0.72. We then repeated this based on the anti-correlated disparity tuning curves (corresponding dashed lines) and the correlation for the example neuron was zero (Figure 4C, dashed red versus dashed light green).

Overall, PV+ disparity tuning to correlated RDS is highly correlated with the average disparity tuning of all surrounding visually responsive PV- neurons within 125 um (Figure 4E, top row). In contrast, disparity tuning for SST+ inhibitory or excitatory neurons have lower correlation with the average disparity tuning of surrounding SST- or other excitatory neurons, respectively (Figure 4D and 4F, top row). This difference is clear across mice, where PV+ inhibitory neuron tuning consistently has a higher correlation with the disparity tuning of surrounding PV- neurons (Figure 4G, red filled circles), as well as in the cumulative distributions of correlation for all neurons grouped together (Figure 4H, solid red line). Because disparity is not uniformly represented in the visual cortex (La Chioma et al., 2019; Brown et al., 2025), there will always be some correlation of disparity tuning between individual neurons and surrounding neurons. Therefore, we also computed what the correlation would be if we shuffled which neurons were compared to which surrounding neurons. This correlation was on average just below 0.16 and is noted by the dashed lines in Figure 4G and 4H. PV+ inhibitory neurons were the only neurons with correlation significantly above this prediction. This suggests that the tuning of PV+ inhibitory neurons is predicted by the tuning of neurons in its local neighborhood well beyond chance and that PV+ inhibitory neurons are integrating the responses of surrounding excitatory neurons relatively non-specifically. The weaker correlation for SST+ inhibitory neurons suggests that they may be integrating the responses of surrounding excitatory neurons in a more selective manner, since the neurons are still as selective for disparity as excitatory neurons (Figure 3G). Since disparity is heterogeneously represented in the visual cortex of mice (Scholl et al., 2015), non-selective integration explains the broader disparity tuning for PV+ neurons (Figure 3D).

Overall, disparity tuning to anti-correlated RDS is not predicted by the activity of surrounding neurons. There is no difference in the correlation between disparity tuning for excitatory neurons and surrounding neurons for correlated RDS, anti-correlated RDS, and the shuffled prediction (Figure 4D, 4G, and 4H). The correlation between disparity tuning for both PV+ and SST+ inhibitory neurons and surrounding neurons though significantly decreases for anti-correlated RDS compared to correlated RDS (Figure 4E-H). This supports that both subclasses of inhibitory neurons are integrating from excitatory neurons in a disparity-dependent manner for correlated RDS (either due to local biases or selective connectivity) and not for anti-correlated RDS as predicted in Figure 2B. The correlation between anti-correlated disparity tuning for PV+ inhibitory neurons and surrounding PV-neurons is not significantly different from excitatory neurons or the shuffled prediction (Figure 4G-H). Although the difference is small, the correlation between anti-correlated disparity tuning for SST+ inhibitory neurons and surrounding SST- neurons is surprisingly significantly lower than what we measure for excitatory neurons or the shuffled prediction (Figure 4H). This is consistent with Figure 3L-M suggesting a large deviation in the energy model prediction for the relationship between correlated and anti-correlated disparity tuning. This provides additional support that SST+ inhibitory neurons might be particularly suited for suppressing false matches in anti-correlated RDS.

### Noise correlation involving PV+ and SST+ inhibitory neurons also suggests different circuitry between subclasses

Correlated variability (noise correlation) between pairs of neurons can also be used to infer connectivity among neurons (Zohary et al., 1994; Shadlen and Newsome, 1998; Smith and Kohn, 2008). If two neurons are connected via a direct or indirect excitatory synaptic connection or receive synaptic inputs from a common input, they will have positive noise correlation. If an increase in activity from one neuron causes a decrease in activity in another neuron due to a direct or indirect inhibitory synaptic connection, they will have negative noise correlation. We measured noise correlation for excitatory-to-excitatory (E-E) neuron pairs, putative excitatory-to-inhibitory (E-I) neuron pairs, and inhibitory-to-inhibitory neuron pairs (within the same subclass) to better understand how different subtypes of inhibitory neurons integrate and modulate disparity information.

Noise correlation has previously been shown to depend on similarity in disparity preference, but not on cortical distance for E-E neuronal pairs (La Chioma et al., 2020). We confirmed that noise correlation for excitatory neuron pairs depended on disparity tuning similarity (signal correlation; Figure 5A, black). Our data suggests that excitatory neurons with more similar disparity tuning are more likely to excite each other (positive noise correlation) and neurons with very different tuning are more likely to suppress each other (negative noise correlation). We found that all combinations of E-I and I-I neuronal pairs also significantly depended on disparity tuning similarity as well (Figure 5A, columns 2-5), which was consistent across mice (Figure 5C). This suggests that there is some organized disparity-dependent circuitry involving both PV+ and SST+ inhibitory neurons. This could be due to local biases in disparity preference (La Chioma et al., 2020; Brown et al., 2025) or due to selective integration of excitatory inputs (Figure 4). This would generate more positive noise correlation among neurons with more similar disparity tuning. Noise correlation does not only represent the excitatory inputs to inhibitory neurons though. It also represents the potential connections from inhibitory neurons back to excitatory neurons that suppress excitatory responses. The negative noise correlation among neuron pairs with very different disparity tuning (negative signal correlation) suggests that both PV+ and SST+ inhibitory neurons could be selectively suppressing non-preferred disparity responses in excitatory neurons (Figure 2F).

**Figure 5.**
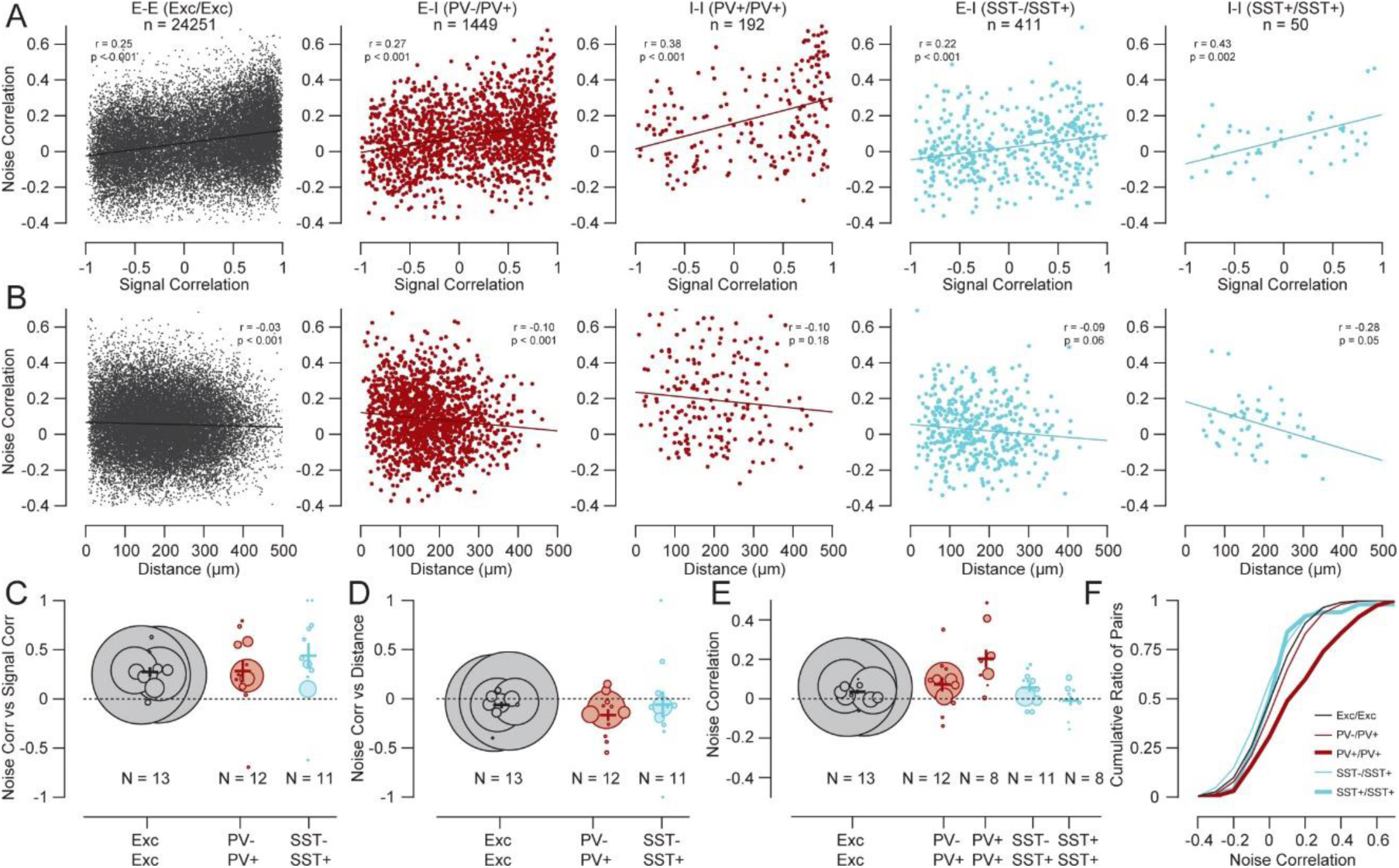
Trial-to-trial covariations are strongest for neuron pairs including PV+ neurons. **A**, For all neuron pairs with significant disparity tuning, trial-to-trial covariation (noise correlation) depends on disparity tuning similarity (signal correlation). **B**, For all neuron pairs with significant disparity tuning, noise correlation depends on distance between neurons. The negative correlation with distance was significantly stronger for PV-/PV+ pairs compared to pairs of excitatory neurons (Steiger Test, *p* = 0.01). **C**, Noise correlation depended on signal correlation consistently across mice (Wilcoxon signed rank test, *p* < 0.001 for Exc/Exc, *p* = 0.02 for PV-/PV+, and *p* = 0.02 for SST-/SST+). **D**, Noise correlation depended on distance only consistently across mice for PV-/PV+ pairs of neurons (Wilcoxon signed rank test, *p* = 0.03). **E**, Based on averages in each mouse, noise correlation is generally small (<0.1) for all pairs of neurons except PV+/PV+ pairs. For PV+/PV+ pairs noise correlation was generally greater than 0.1 and was significantly higher compared to pairs of excitatory neurons in mice (Wilcoxon rank sum test, *p* = 0.004). **F**, The aggregate distribution of noise correlation in all PV+/PV+ neurons is clearly and significantly larger than the noise correlation measured in all other pairs of neurons not involving PV+ neurons (Wilcoxon rank sum test, *p* < 0.001).

We observed that the dependence of noise correlation on cortical distance for excitatory neurons was very weak, but was still statistically significant (Figure 5B, black). The dependence of noise correlation on cortical distance appeared to be stronger for neuronal pairs involving inhibitory neurons (E-I or I-I) compared to E-E pairs although the relationship was only statistically significant for PV+-to-PV- pairs of neurons (Figure 5B and 5D). This suggests that circuits involving PV+ inhibitory neurons in particular are localized compared to other circuits.

The most contrasting noise correlation result we found between excitatory neurons and subclasses of inhibitory neurons was that noise correlation was higher when PV+ neurons were part of the pair (Figure 5E-F). Noise correlation is generally on average much smaller than 0.1, but the average noise correlation among PV+ neuronal pairs was 0.2. This higher noise correlation is consistent with PV+ inhibitory neurons integrating inputs from excitatory neurons less discriminately compared to SST+ inhibitory neurons. Greater numbers of connections between PV+ inhibitory neurons and excitatory neurons would increase their shared variability and increase the amount of shared input among pairs of PV+ to PV+ inhibitory neurons.

Our measures of functional connectivity by noise correlation and the spatial integration of PV+ and SST+ neurons indicate differences in the network connectivity of these neurons relative to their excitatory counterparts. PV+ neurons appear to be more coupled to nearby excitatory cells and to their own population than any circuits involving SST+ neurons. This dense connectivity by PV+ cells likely underlies their weak disparity selectivity to correlated RDS. In contrast, SST+ neurons display noise correlation structures and spatial integrative properties that appear more selective, akin to their excitatory counterparts. This specificity, and the lack of a 180-degree inversion for anti-correlated RDS, suggest that they may play a role in suppressing false matches between the two eyes.

### PV+ and SST+ inhibitory neurons sharpen disparity tuning to correlated RDS

A limitation of noise correlation (Figure 5) is that the interaction between two neurons is only represented by a single value. Interactions between excitatory and PV+ or SST+ inhibitory neurons are in both directions. Our measurements of correlation between disparity tuning of PV+ or SST+ neurons and disparity tuning of the average activity of surrounding PV- and SST- neurons focus on the direction from excitatory to inhibitory neurons (Figure 4). These results revealed to us the organization of how PV+ and SST+ inhibitory neurons integrate the responses of excitatory neurons. To focus on the other direction, we needed to measure the influence of these inhibitory cell classes on disparity selectivity in excitatory neurons. We employed optogenetics to selectively suppress the activity of either PV+ or SST+ neurons and then measured how this affected disparity tuning to correlated and anti-correlated RDS of the remaining population. Our goal was to suppress inhibitory responses, but not to inactivate these neurons completely. If either of these cell classes plays a role in sharpening selectivity for disparity, then suppressing them should reduce disparity selectivity to correlated RDS. On the other hand, if these cell classes play a role in suppressing false matches present in anti-correlated RDS, then we expect anti-correlated selectivity to increase with optogenetic suppression.

PV-Cre and SST-Cre animals were injected with an adeno-associated virus labeling all neurons with GCaMP6s and only neurons expressing Cre (PV+ or SST+) with TdTomato and Archaerhodopsin-3 (Arch). Arch is a light-activated outward proton pump that hyperpolarizes the cell when triggered by green-yellow light and optogenetically inhibits neuronal responses (Chow et al., 2010). We used ArchT, which is an Arch variant with increased light sensitivity and can inhibit spiking in neurons with 595 nm (orange) light. We used an LED delivering 590 nm light routed through the objective every other frame to quasi-simultaneously induce optogenetic suppression of PV+ or SST+ neurons while imaging calcium responses from all neurons (Figure 6A, right). In order to reduce any light from the LED inadvertently being recorded as signal by the PMT, we added a pi cell liquid crystal shutter in front of the PMT (Figure 6A, left). The LED was synchronized with the pi cell and only delivered light when the shutter was closed (Figure 6A, dashed line). We only analyzed frames in which the shutter was open and the LED was off. LED power was calibrated by measuring it at the objective and testing where the responses of PV+ and SST+ neurons had their responses reduced to 50% on average (Figure 6B-C). Since the red channel was used to deliver the LED light to the window during two-photon imaging, we used changes in the response properties to LED activation to classify neurons as PV+ versus PV- and SST+ versus SST-. Neurons with a significant decrease in response in the 2-s period (30 frames) when the LED was on compared to the response when the LED was off before stimulus onset were classified as PV+ and SST+ inhibitory neurons. All other neurons were classified as PV- and SST-. PV+ and SST+ inhibitory neurons were consistently suppressed by the LED during visual evoked activity, as well (Figure S2, red). Visually evoked responses for SST- neurons were generally disinhibited by SST+ suppression, but responses for PV- neurons were not clearly disinhibited or suppressed by PV+ suppression (Figure S2, blue).

**Figure 6.**
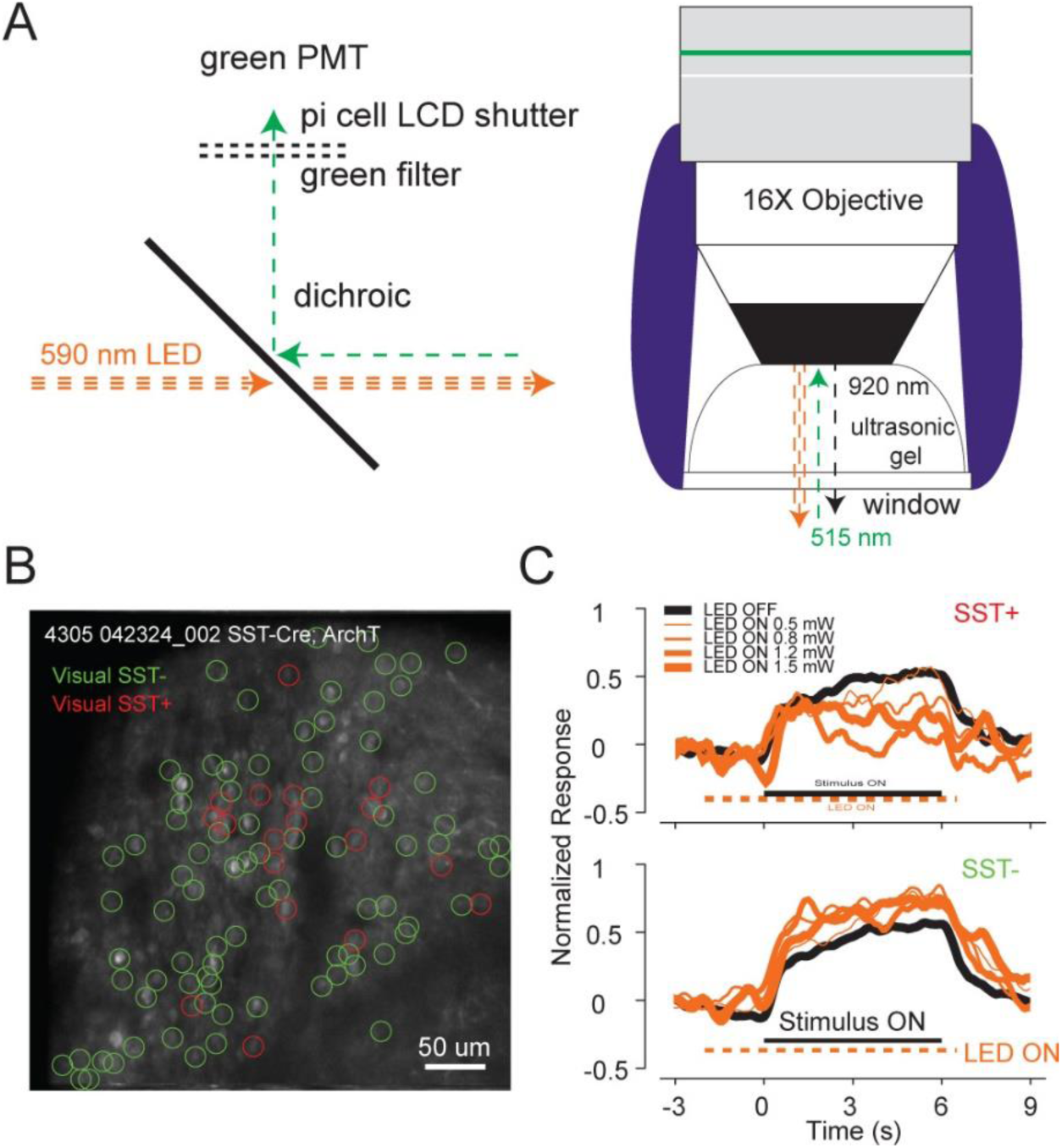
Optogenetics experimental setup. **A**, Left, we used a pi cell liquid crystal shutter in front of the PMT to reduce light bleedthrough. The LED was also synchronized to the pi cell and only delivered light when the shutter was closed (dashed orange line). Right, for even illumination of the imaging field, the LED light was routed through the objective. **B**, Example SST-Cre animal with ArchT. Red circles identify visual SST+ neurons and green circles identify visual SST- neurons. **C**, We varied the LED power and measured the normalized response in both SST+ and SST- neurons (the same neurons in Figure 5B). We found that the SST+ neurons had a decrease in activity when the LED came on, and the SST- neurons had an increase in activity, confirming that the optogenetic manipulation was working.

We first examined how disparity tuning changed in putative excitatory neurons in response to correlated RDS with optogenetic suppression of inhibitory neuron responses. We found in general that disparity tuning broadens after PV+ or SST+ suppression (Figure 7A and 7F) with consistently reduced amplitude modulation (Figure 7B and 7G). The broader tuning and reduced amplitude modulation of PV- and SST- neurons with PV+ and SST+ suppression, respectively, was generally associated with stronger disinhibition at the least preferred disparities (Figure 7A and 7F). Overall, these data show that PV+ and SST+ inhibitory subclasses of neurons sharpen disparity tuning in excitatory neurons by suppressing the responses to non-preferred disparities consistent with our prediction in Figure 2F.

**Figure 7.**
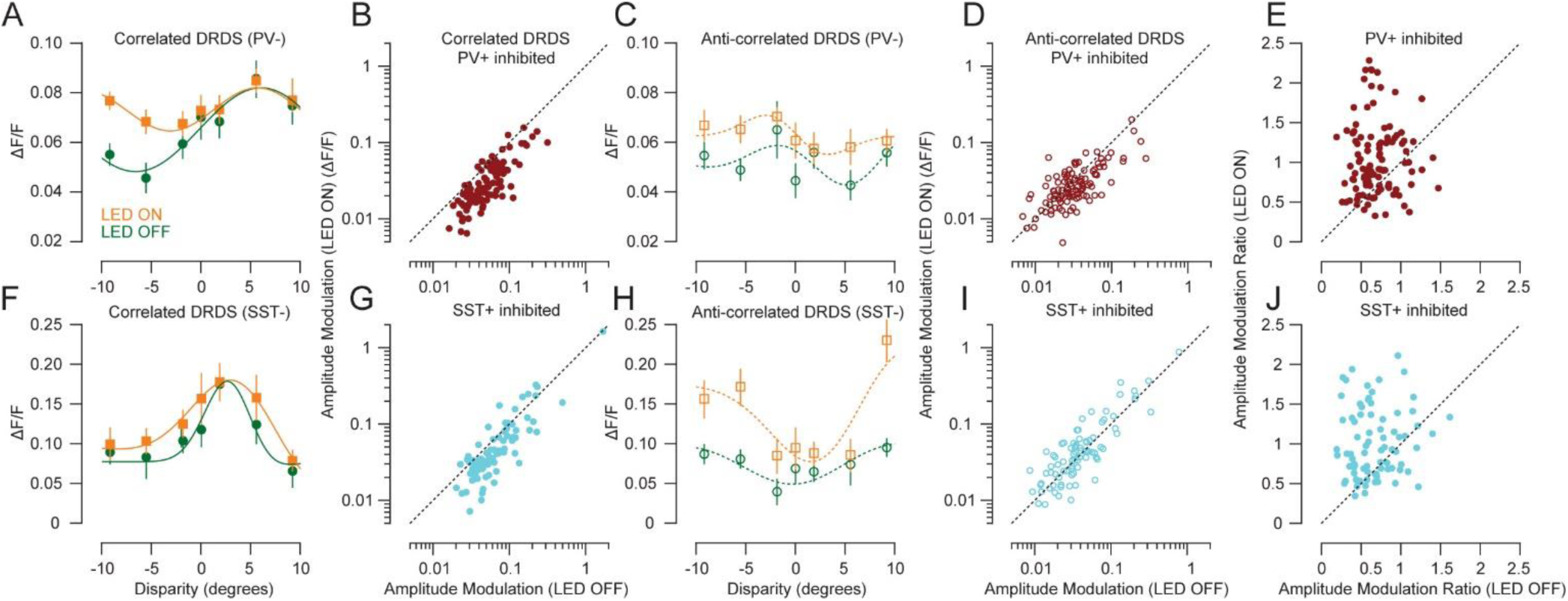
Suppression of PV+ and SST+ inhibitory neurons increases the amplitude modulation ratio. **A**, Optogenetic suppression of PV+ inhibitory neurons broadens disparity tuning to correlated RDS for PV- neurons. Error bars are standard error (n = 10 trials). **B**, The amplitude modulation for disparity tuning to correlated stereograms significantly decreased for PV- neurons during optogenetic suppression of PV+ inhibitory neurons (Wilcoxon signed rank test, p < 0.001). **C**, Optogenetic suppression of PV+ inhibitory neurons also broadens disparity tuning to anti-correlated RDS for PV- neurons (same neuron as A). **D**, The amplitude modulation for disparity tuning to anti-correlated stereograms also significantly decreased for PV- neurons during optogenetic suppression of PV+ inhibitory neurons (Wilcoxon signed rank test, p < 0.001). **E**, The amplitude modulation ratio between anti-correlated and correlated disparity tuning significantly increased to close to 1 (from 0.65 to 0.92) when PV+ neurons were optogenetically suppressed (n = 111 neurons, Wilcoxon signed rank test, p < 0.001). There was no significant change in the correlation between correlated and anti-correlated disparity tuning (Wilcoxon signed rank test, p = 0.83). **F**, Optogenetic suppression of SST+ inhibitory neurons broadens disparity tuning to correlated RDS for SST- neurons. **G**, The amplitude modulation for disparity tuning to correlated stereograms significantly decreased for SST- neurons during optogenetic suppression of SST+ inhibitory neurons (Wilcoxon signed rank test, p < 0.001). **H**, Optogenetic suppression of SST+ inhibitory neurons sharpens disparity tuning to anti-correlated RDS for SST- neurons (same neuron as F). **I**, The amplitude modulation for disparity tuning to anti-correlated stereograms significantly increased for SST- neurons during optogenetic suppression of SST+ inhibitory neurons (Wilcoxon signed rank test, p = 0.03). **J**, The amplitude modulation ratio between anti-correlated and correlated disparity tuning significantly increased to close to 1 (from 0.59 to 0.92) when SST+ neurons were optogenetically suppressed (n = 84 neurons, Wilcoxon signed rank test, p < 0.001). There was no significant change in the correlation between correlated and anti-correlated disparity tuning (Wilcoxon signed rank test, p = 0.52).

### PV+ inhibitory neurons sharpen and SST+ inhibitory neurons broaden disparity tuning to anti-correlated RDS

Similar to excitatory neurons (Figure 1C-D), PV- and SST- neurons generally responded to anti-correlated RDS with an inversion in tuning of a reduced magnitude compared with the response to correlated RDS (e.g., Figure 7C and 7H versus Figure 7A and 7F). We hypothesized that inhibitory neurons could cause this reduced magnitude in response to anti-correlated RDS. PV+ or SST+ could be selectively inhibiting the responses to false matches, thus helping to solve the stereo correspondence problem. If this were the case, when PV+ or SST+ neurons are optogenetically suppressed, we should expect to see larger amplitude modulation in disparity tuning to anti-correlated RDS.

For PV- neurons, suppression of PV+ inhibitory responses broadened disparity tuning to anti-correlated RDS in a similar manner as observed for disparity tuning to correlated RDS (Figure 7C). The amplitude modulation of PV- neurons was consistently reduced by suppression of PV+ inhibitory neurons (Figure 7D). The reduction in amplitude modulation for disparity tuning to anti-correlated RDS caused by PV+ suppression though was significantly smaller than the reduction in amplitude modulation for disparity tuning to correlated RDS (Wilcoxon signed rank test, p < 0.001). This produces an increase in amplitude modulation ratios with suppression of PV+ inhibitory neurons (Figure 7E). Therefore, although PV+ inhibitory neurons did not selectively suppress false matches, they more effectively sharpened disparity tuning for correlated than anti-correlated RDS.

For a majority of SST- neurons, suppression of SST+ inhibitory responses sharpened disparity tuning to anti-correlated RDS (Figure 7H). Although the effect was modest on average, the amplitude modulation of SST- neurons significantly increased from an average of 0.06 to 0.07 with suppression of SST+ inhibitory neurons (Figure 7I). The combination of an increase in amplitude modulation for disparity tuning to anti-correlated RDS and a reduction in amplitude modulation for disparity tuning to correlated RDS with suppression of SST+ inhibitory neurons results in an increase in amplitude modulation ratios (Figure 7J).

## Discussion

The disparity energy model predicts that the 180-degree phase-shifted disparity tuning to anti-correlated RDS should have the same magnitude of amplitude modulation as disparity tuning to correlated RDS (Ohzawa et al., 1990). We have shown that disparity tuning to anti-correlated stereograms in excitatory neurons has weaker modulation with respect to correlated stereograms (Figure 1), in agreement with previous work (Cumming and Parker, 1997; Ohzawa et al., 1997; La Chioma et al., 2020). We hypothesized that circuits involving PV+ and SST+ inhibitory neurons could suppress the false matches found in anti-correlated RDS and explain their reduced amplitude modulation (Figure 2). Our optogenetic results suggest that PV+ inhibitory neurons sharpen disparity tuning to correlated RDS and sharpen disparity tuning to anti-correlated RDS to a lesser extent for excitatory neurons (Figure 7C-D). We found that SST+ neurons sharpen disparity tuning to correlated RDS and had a small, but significant effect consistent with SST+ inhibitory neurons broadening disparity tuning to anti-correlated RDS for excitatory neurons (Figure 7H-I). This supports that the SST+ microcircuit selectively suppresses false matches and indicates that PV+ and SST+ inhibitory microcircuits perform distinct computations with respect to stereo matching.

The microcircuit involving PV+ inhibitory neurons has characteristics that are consistent with previous work (Scholl et al., 2015) and consistent with our prediction in Figure 2C. Multiple analyses indicated that PV+ neurons broadly pool local disparity information, including less selective disparity tuning than neighboring excitatory neurons, stronger correlation with local population responses, and higher noise correlations. These observations support the idea that PV+ neurons receive substantial shared input and regulate the gain of local excitatory populations rather than performing highly selective stereo computations (Atallah et al., 2012; Wilson et al., 2012). Nonetheless, optogenetic suppression of PV+ neurons broadened disparity tuning in excitatory neurons, indicating that PV+ inhibition sharpens disparity selectivity. This effect is likely possible because PV+ neurons are themselves mildly disparity-tuned and because local biases in disparity preference exist within mouse visual cortex (La Chioma et al., 2019; Brown et al., 2025). Notably, the sharpening produced by PV+ neurons was stronger for correlated than anti-correlated RDS. Together with the weaker agreement between PV+ responses and disparity energy model predictions for anti-correlated stimuli, these results suggest that PV+ neurons enhance responses to correct matches more effectively than responses to false matches. Thus, while PV+ neurons do not appear to selectively suppress false matches, they may contribute to stereo correspondence by improving the representation of correct matches.

The microcircuit involving SST+ inhibitory neurons exhibited several properties distinct from the PV+ inhibitory microcircuit and consistent with the hypothesis in Figure 2D. In contrast to PV+ neurons, SST+ inhibitory neurons showed evidence of more selective integration of disparity information, including weaker shared variability with neighboring neurons and tuning that was less predictable from the surrounding population. Optogenetic suppression of SST+ inhibitory neurons broadened disparity tuning indicating that both inhibitory cell classes sharpen selectivity (Figure 7F-G). SST+ neurons, however, deviated substantially from disparity energy model predictions and was poorly correlated with surrounding neurons (even below the shuffled prediction), suggesting specialized processing of disparity information. Consistent with this idea, SST+ inhibitory neurons broadened selectivity to anti-correlated RDS, demonstrating a role for suppressing false matches. An alternative possibility is that the disparity tuning for anti-correlated RDS is on average relatively flat (e.g., Figure 3I, right) in SST+ inhibitory neurons and their suppression of excitatory responses is pulling responses below the spiking threshold (Figure S2) (Xu et al., 2026). Regardless of the underlying mechanism, SST+ inhibitory neurons appeared to be especially suited to play an important role in stereo matching by both enhancing correct matches and suppressing false matches.

There are a few major limitations of our data that are important to our interpretations. First, we were not ever simultaneously observing excitatory, PV+ inhibitory, and SST+ inhibitory neurons at the same time. We were measuring either excitatory neurons alone, or PV+ inhibitory neurons compared to all other neurons (PV-, excitatory and other inhibitory neurons), or SST+ inhibitory neurons compared to all other neurons (SST-, excitatory and other inhibitory neurons). Even though the majority of “all other neurons” will be excitatory neurons, we cannot rule out that other inhibitory subclasses contributed to any effects that we observed. Inhibitory neurons with high spiking rates might even have a larger representation in our optogenetic experiments because these cells have brighter GCaMP responses. The pi cell we used is polarized introducing additional filtering that could mask excitatory neurons that are relatively less responsive with lower brightness. Second, inhibitory neurons also receive inputs from other inhibitory neurons. Inhibitory inputs to PV+ inhibitory neurons come mostly from layer 2/3, whereas SST+ inhibitory neurons receive inhibition from layers 2/3, 4, and 5 (Xu and Callaway, 2009). Recent work has shown that SST+ inhibitory neurons inhibit PV+ inhibitory neurons (Cottam et al., 2013; Pfeffer et al., 2013; Pi et al., 2013) and PV+ neurons also inhibit SST+ neurons, albeit less strongly (Pfeffer et al., 2013; Walker et al., 2016). Because SST+ inhibitory neurons are known to strongly inhibit PV+ inhibitory neurons, some of the sharpening of disparity tuning to anti-correlated RDS with suppression of SST+ inhibitory neurons could be due to PV+ inhibitory neuron disinhibition (Cardin, 2018). This seems unlikely though since PV+ inhibitory neurons sharpen disparity tuning more strongly for correlated RDS compared to anti-correlated RDS (Figure 7B and 7D) and we found that disparity tuning to correlated RDS actually broadened with suppression of SST+ inhibitory neurons. Lastly, even with SST+ suppression of false matches, excitatory neurons in the visual cortex still have disparity-dependent responses to anti-correlated RDS so additional mechanisms that were not examined in this study are necessary to solve the stereo correspondence problem completely.

The results of our experiments provide important constraints as to how distinct cortical circuits contribute to solving the stereo correspondence problem. We revealed that PV+ and SST+ inhibitory neurons are likely enhancing selectivity to correlated RDS and effectively amplifying correct matches. The anti-correlated optogenetic results reveal differences between these subclasses of inhibitory neurons that are consistent with differences in their disparity tuning properties and organization within the cortical network. This distinction supports the conclusion that SST+ inhibitory neurons play an additional important role in suppressing the responses to false matches.

## Acknowledgments

We are grateful to Boris Zemelman for providing the equipment needed and viruses necessary to carry out the optogenetic experiments. We thank Stefanie Esmond, Ronan O’Shea, Vy Nguyen, and Mason Antin for providing technical assistance.

There are no conflicts of interest.

