## Supplementary material for "Parvalbumin and somatostatin inhibitory microcircuits perform distinct roles in stereo matching": Figure S1

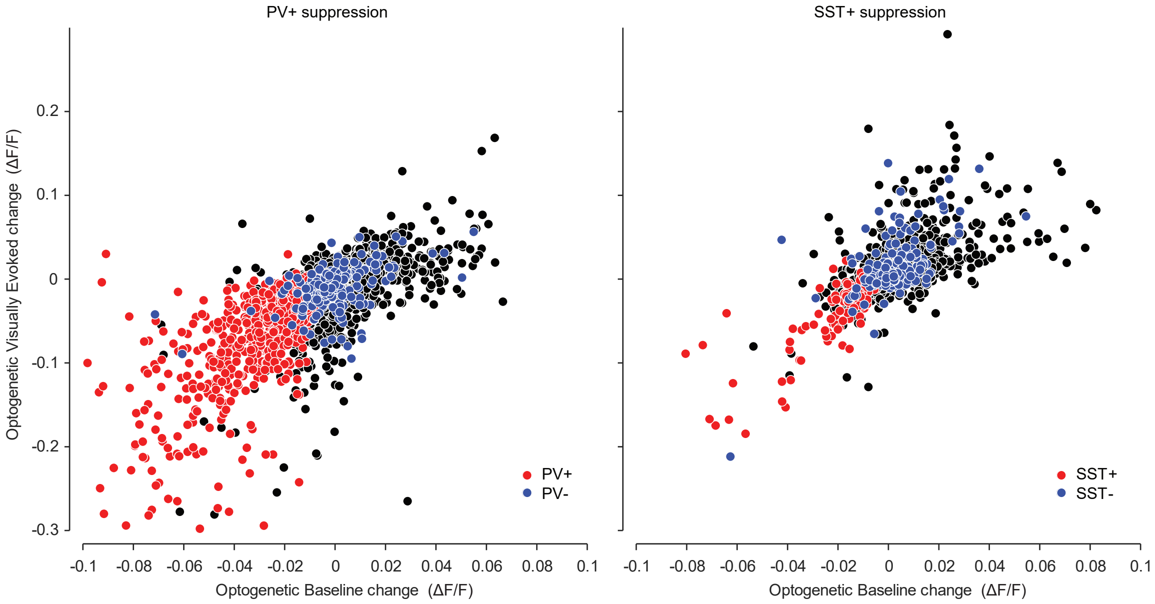


**Figure S2. ArchT inhibits PV+ and SST+ cells and disinhibits PV- and SST- cells.**
Both plots show all visually responsive cells from two experiments, one with PV+ cell suppression (left) and one with SST+ cell suppression (right). Red circles indicate PV+ and SST+ cells, blue circles indicate tuned PV- or SST- cells, and black circles indicate untuned PV- or SST- cells. Untuned PV- or SST- cells were not included in further analysis.
