## Supplementary material for "Parvalbumin and somatostatin inhibitory microcircuits perform distinct roles in stereo matching": Figure S2

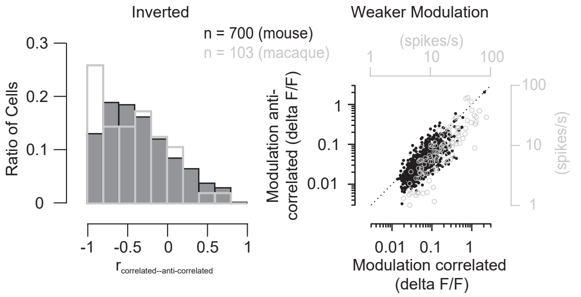


**Figure S1. Disparity tuning to anti-correlated stereograms is inverted and has weaker modulation with respect to correlated stereograms similar to macaque V1.**
Left, The correlation between corr and anticorr disparity tuning for mice was similar to what is observed in macaques (Samonds et al. 2013). Right, The modulation (maximum-minimum response) for anticorr disparity tuning for mice was similar to what is observed in macaques (Samonds et al. 2013).
